# QUILPEN provides independent and label-free single-cell quantification of pigmentation dynamics and organelle content

**DOI:** 10.1101/2025.10.20.683473

**Authors:** Rebecca G. Zitnay, Shukran Alizada, Tarek E. Moustafa, Devin Lange, Luke Schreiber, Alexander Lex, Robert L. Judson-Torres, Thomas A. Zangle, Rachel L. Belote

## Abstract

The relationship between melanogenesis, pigmentation, and melanocyte behavior is complex. In melanocytes, pigmentation is often associated with differentiation, yet mature melanocytes vary in pigment content. In melanoma, pigmentation-linked transcriptional programs may have prognostic value, but visual assessments of tumor pigmentation have yielded inconsistent results. Progress linking pigmentation phenotypes to cell state has been limited by a lack of tools that can directly and dynamically quantify melanin content in live cells. Here we present QUantitative Imaging of Label-free Pigment-associated ENtities (QUILPEN), a label-free multi-modal imaging technique that combines quantitative phase imaging (QPI), quadrant darkfield (QDF), and absorption imaging, to independently capture light that has been transmitted, scattered, and absorbed. This non-destructive method enables live-cell imaging over multiple days without labels. We show absorption as a reliable readout of melanin content, which can be decoupled from melanosome content detected by QDF, which measures scattered light. Applying QUILPEN to melanoma cells before and during repigmentation, we find that melanin content is highly heterogeneous, and that this heterogeneity is reinstated upon repigmentation. Lineage tracking further reveals that melanin synthesis rates are heritable and can be transmitted both symmetrically and asymmetrically. QUILPEN enables real-time quantification of pigmentation dynamics and cell-level heterogeneity.

**Significance:** Pigmentation is a defining feature of melanocytic cells and linked to behavior, yet melanin’s optical properties complicate direct measurement with existing approaches. Taking advantage of the absorbing nature of pigment and scattering properties of organelles, we can independently track pigment and organelle content over time and along cell lineages in a label-free manner to reveal population heterogeneity and patterns of inheritance.

**Research Highlights:** QUILPEN is a label-free, live cell imaging pipeline that independently quantifies pigment and organelles in melanocytic cells. Tracking pigment and organelle content along cell lineages reveals varied inheritance of pigment-associated features.

**Graphical Abstract:** 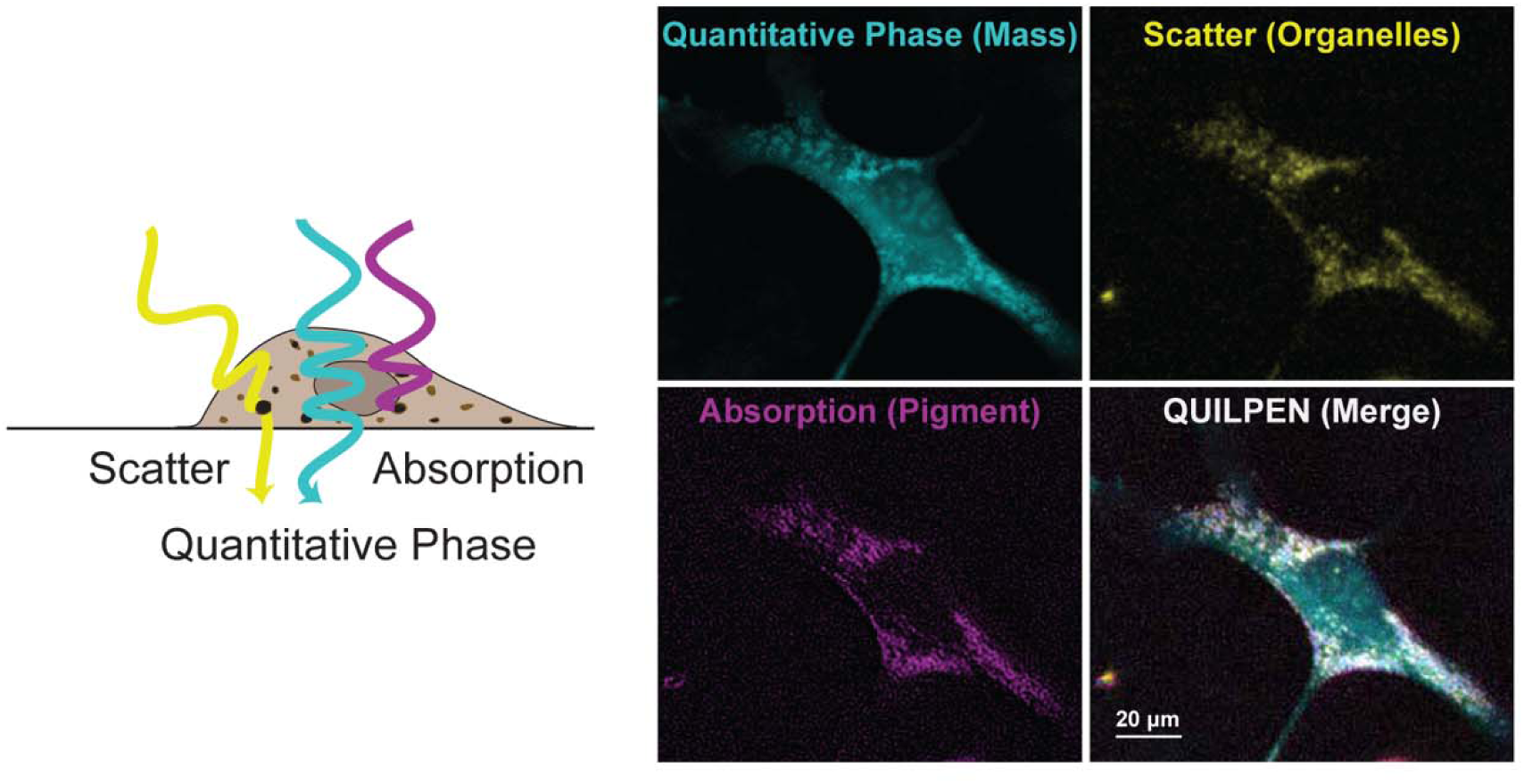

## Introduction

Melanocytic cells are distinguished by their ability to synthesize, package, and distribute melanin, a multifunctional pigment involved in UV protection, oxidative stress regulation, and metal ion homeostasis. The pigmentation phenotype is associated with specific cellular behaviors and developmental programs. Pigment-associated gene expression has been used as a proxy for melanocyte differentiation, proliferation, and metabolic status (Bajpai et al., 2023; Huberman et al., 1979; Mica et al., 2013). In melanoma, pigmentation-associated gene expression correlates with prognosis (Akbani et al., 2015; Lourenço et al., 2020; Netanely et al., 2021) and pigment itself influences both immune recognition and therapeutic response (Cabaço et al., 2022; Chen et al., 2009; Mohagheghpour et al., 2000). However, pigment content and transcriptional programs do not always align. For example, pigmentation is often considered a marker of differentiated cell state although single-cell studies have shown that mature melanocytes can vary widely in both pigment content and expression of pigment-associated genes (Belote et al., 2021). In the context of oncogenic transformation, melanoma cells also exhibit heterogeneity in both pigmentation and expression of pigment-associated genes (Fedele et al., 2025; Rambow et al., 2018). In short, the relationships between pigment content, transcriptional programs, and cell state observed in bulk population studies become markedly more complex when examined at single-cell resolution. This complexity highlights the need for tools that can directly measure pigment in live cells while capturing associated cellular features over time.

Melanogenesis is a dynamic and complex process involving hundreds of genes (Bajpai et al., 2023; Baxter et al., 2019). Current understanding of melanin synthesis, melanosome biogenesis, and pigment function derives largely from bulk analysis conducted at discrete time points using purified melanin, isolated melanosomes, or fixed cells. Quantification of melanin by spectroscopy, mass spectrometry, or high-performance liquid chromatography (HPLC) has yielded valuable insights into pigment chemistry, optical properties, and relative abundance of melanin within cells (Ito et al., 2011; Sardar et al., 2001). Isolation and fractionation of bulk melanosomes have identified stage-specific protein composition (Chi et al., 2006; Hoashi et al., 2005; Kushimoto et al., 2001). Imaging methods, such as Fontana Masson, immunofluorescence, and electron microscopy, have yielded spatial and structural details about melanosome maturation and distribution (Ando et al., 2011; Hoashi et al., 2005; Peles et al., 2009). However, each of these methods is inherently destructive and requires labeling or fixation and, therefore, does not permit live-cell analysis. The primary non-destructive technique for assaying the pigmentation phenotype is flow cytometry side scatter, which correlates with intracellular granularity and tends to be high in darker pigmented cells or cells that have internalized melanosomes (Belote et al., 2021; Choi et al., 2012). However, this approach cannot distinguish melanin content from other cellular features that influence granularity, such as melanosome content, nor does it permit dynamic tracking in living cells.

Quantifying melanin in individual living cells is technically challenging due to the optical properties of the pigment. Melanin strongly absorbs and scatters light across a broad spectrum (Riesz et al., 2006), leading to reduced excitation and fluorescence quenching that interferes with conventional light-based readouts (Stockert et al., 2022). These properties hinder quantitative assessment of pigment content and complicate the use of fluorescent reporters or dyes in pigmented cells. Melanosome-targeted biosensors can report on organelle localization or pH, but require fluorescent labels that may be quenched by melanin and do not provide information about pigment content (Bruder et al., 2012; Scales et al., 2021). Autofluorescence lifetime imaging can detect endogenous melanin and report on aspects of the local biochemical environment. However, it does not provide a direct or quantitative measure of melanin and is constrained by spectral overlap with other endogenous fluorophores and the requirement for high-intensity pulsed laser illumination (Krasieva et al., 2012; Pena et al., 2022; Sitiwin et al., 2019).

Here, we introduce a label-free, non-destructive, low-phototoxicity imaging approach that enables time-resolved, single-cell analysis of melanin and associated cellular features in live melanocytic cells. Our approach, QUantitative Imaging of Label-free Pigment-associated ENtities (QUILPEN) is built on a custom multi-modal quantitative phase imaging microscope that independently measures transmitted, scattered, and absorbed light, yielding high-content biophysical profiles of live melanocytic cells. We use this system to quantify pigment heterogeneity across melanoma cell populations, decouple melanin content from organelle content, and track pigmentation dynamics over time. Integration with open-source tracking (Ershov et al., 2022) and visualization (Lange et al., 2022, 2025) tools enable lineage tracing and support investigation of how pigmentation state relates to phenotype inheritance across generations. Together, these capabilities establish QUILPEN as a versatile platform for investigating melanogenesis and pigment-associated phenotypes in healthy and disease contexts.

## Materials and Methods

### Cell culture

Cells were passaged 2-3 times a week to maintain a confluence between 30 and 90% and maintained at 5% CO_2_ at 37°C. Cells were passed by washing with DPBS (#14190250, Thermo Fisher Scientific) and dissociated with Trypsin-EDTA (#25300120, Thermo Fisher Scientific) or TrypLE (#12605010, Thermo Fisher Scientific). MNT-1 cells (ATCC, CRL-3450) were cultured in Dulbecco’s Modified Eagle’s Medium (DMEM; ATCC 30-2002) with 20% FBS (VWR, 89510-186), and NEAA (1%, ThermoFisher Scientific, 11140050). SK-MEL-28 cells (CVCL_8054, ATCC) and were cultured in RPMI 1640 with GlutaMax (#61870036, Thermo Fisher Scientific) supplemented with 10% FBS (VWR, 89510-186). Patient-derived xenograft lines of darkly pigmented acral melanoma (AM084, formerly MTG084) and lightly pigmented subungual acral melanoma (ASM021, formerly MTG021) were previously generated (Moustafa et al., 2024; Smith et al., 2024) and grown in AM3 media (80% MCDB153 (#M7403, Sigma, United States), 20% L-15 media (#11415-064, Thermo Fisher Scientific), 2.5% FBS (#FB5001-H, Thermo Fisher Scientific), 1X Insulin-Transferrin-Selenium X (#51500-056, Thermo Fisher Scientific), 5 ng⁄mL EGF, 0.2% BPE, 10 ng⁄mL insulin-like growth factor, 5 μg⁄mL transferrin, 3 ng⁄mL BFGF, 3 μg⁄mL heparin, 0.18 μg⁄mL hydrocortisone, 10 nM endothelin 1, and 1.68 mM CaCl2). Penicillin-Streptomycin (1%, 15-140-122, Thermo Fisher Scientific) was added 24-48 hours before and during imaging. Cells were plated in a 24 well dish (662160, Greiner Bio-One, Germany) at a density of 5000 to 8000 cells per well 40 hours prior to imaging

### PTU Treatment

N-Phenylthiourea (PTU, P7629, Sigma-Aldrich) was prepared at 0.1 M in DMSO. PTU was added to the cells for a final concentration of 600 µM in DMSO (0.6%). Cells were cultured in PTU for at least 21 days before use in experiments.

### Protocol for multi-channel Image Acquisition and Tracking Analysis

#### Protocol A, 3-Channel Image Acquisition and Processing (Supplementary Fig. 1A)

1. Microscope Construction
  a. To perform QPI, Absorption, and QDF imaging construct the LED-based microscope and place it inside a tissue culture humidified incubator to maintain the cells at 37°C and 5% CO_2_ (Parts list and assembly instructions available (Moustafa et al., 2023). Here imaging was performed with a 0.25 NA, 10× objective (PLN 10×, Olympus, Japan) and a monochrome 1920 × 1200 CMOS camera (GS3-U3-23S6M-C, Teledyne FLIR, United States) with an illumination coherence factor of 1.25 (Tian & Waller, 2015).
  b. (Optional) To reduce background artifacts from the LED array, mount a 220-grit ground glass diffuser (47-951, Edmund Optics) in the light path abutting the LED array. This can be performed using hot glue or with a custom-designed, 3D-printed holder.
2. Acquire Images
  a. Follow protocol previously reported for combined QPI and QDF imaging (Moustafa et al., 2024)
    i. Imaging position selection. To avoid scattering artifacts from the edge of the well, imaging locations should be limited to 3 x 3 grid of 9 positions at the center of the well.
    ii. At each imaging location, capture a total of 8 images with different LED illumination patterns: four images for differential phase contrast QPI (right half, left half, top, and bottom), and four images for QDF (top-right, bottom-right, top-left, and bottom-left)
    iii. Specify imaging dynamics. We capture 360 frames over 120 hours with a 20-minute time step and single autofocus between cycles.
3. Process raw images
  a. Compute differential phase contrast (DPC) by Tikhonov regularization (Tian et al., 2014; Tian & Waller, 2015). Process the resulting phase and absorption images using custom Matlab scripts (https://github.com/Waller-Lab/DPC/, https://github.com/Zangle-Lab/MultiparametricQPI, and https://github.com/Zangle-Lab/QDF described previously (Moustafa et al., 2024; Polanco et al., 2022; Tian & Waller, 2015).
    i. For DPC, perform background correction by masking cells, averaging background pixels to obtain a common background image, subtracting the common background image, and finally fitting and removing an eighth-order polynomial to the remaining background signal.
    ii. QDF background correction was performed by subtracting an image of an empty reference field prior to segmentation of cells and fitting the residual background signal to an eighth-order polynomial (Moustafa et al., 2024)
4. Matlab output to TIFF images
  a. Matlab output of microscope code will return a .m file for each image position. Run *LoopFolder_quilExportTiffs.m* for positions that pass quality control (no major artifacts, good seeding density). The output folder contains TIFF files of selected images. Absorption images are inverted during processing so that a higher pixel value (lighter color) corresponds to a greater magnitude of absorption

#### Protocol B, Image analysis for feature and lineage tracking (Supplementary Fig. 1B)

1. High fidelity analysis (Single frame analysis: Fig. 2, 3; Time course: Fig. 4F-G, 5)

a. Export selected frame(s) using ImageJ (Schindelin et al., 2012).

i. Select artifact-free frames that have appropriate cell density for segmentation.
b. Segment cells with Cellpose-SAM

i. Install and run Cellpose-SAM using gpu in the python graphical user interface (GUI) using a python virtual environment. (Pachitariu et al., 2025; Pachitariu & Stringer, 2022)
ii. Manually correct Cellpose segmentation using draw, merge, and delete tools in the Cellpose interface.
iii. Use corrected images to train a human-in-the loop model (Pachitariu et al., 2025). Corrected model is helpful to minimize correction time in longitudinal data sets and will be used by the TrackMate-Cellpose plugin.
c. Record measurements (single frame)

i. Export ImageJ ROIs in Cellpose GUI
ii. For measurements of objects within a single frame, import the ImageJ ROIs into ImageJ from cellpose and images from the three image modalities (phase, QDF, absorption). Record integrated density measurements for all segmented objects using *measure* in the ROI window. Export the resulting CSV file.
d. TrackMate with label image (time course)

i. Export label image from Cellpose.
ii. Prepare a four-channel image by merging the phase, absorption, QDF, and label image as separate channels in ImageJ.
iii. Use TrackMate ImageJ plugin (Ershov et al., 2022) with the label detector and set tracking parameters based on cell size and behaviour. For MNT1s, we applied a 50.0 µm max distance for frame-to-frame linking, a 25.0 µm max distance for segment gap closing and allowed for track segment splitting with a max distance of 55.0 µm.
iv. Manually correct all tracks using *trackscheme* feature.
v. Rename tracks in TrackMate GUI by appending “a” or “b” after cell divisions. For the high-fidelity tracking, all tracks were manually corrected. The “spots” feature table and ImageJ ROIs were exported as a CSV file for further analysis in Matlab and Graphpad PRISM.
2. Bulk cell analysis

a. Prepare 3-channel image stack

i. Prepare 3-channel multi-TIFF image stack from TIFF time stacks for each channel (Phase, Absorption, QDF).
ii. Check image Properties to ensure the correct x, y, z dimensions (ie. Make sure frames and slices are not swapped)
b. Segment using TrackMate-Cellpose

i. Download ImageJ plugin for TrackMate-Cellpose and set the appropriate directory for the Cellpose Installation location.
ii. Open the TrackMate-Cellpose plugin, select Cellpose detector and call the path to the Cellpose model that best matches your dataset trained in 1.b.iii.
iii. Run the plugin, selecting the parameters that best match your dataset.
iv. Export spots table as a .CSV file.
3. Track post-processing and plotting

a. Convert data for plotting

i. Run *UnstackedAnalysisOfQUILData.m* (Matlab) to read in trackmate CSV and organize data by tracks and generate track summary information for analysis in graphpad PRISM.
b. Prepare plots

i. (optional) Run *AppendTablesFromDirectories.m* to combine ROI properties exported from CSVs from FIJI stored in subdirectories for each image
ii. (optional) *TimeBin_SplitViolin.m* can be used to prepare the split violin for fig. 4
iii. Final analysis and plotting in GraphPad PRISM
4. TrackMate to Loon

a. Reformat TrackMate output for Loon

i. Detailed instructions for preparing TrackMate files and running the script are available at https://vdl.sci.utah.edu/loonar/docs/getting-started-with-loon/trackmate-data/. Briefly,
ii. Prepare Input: Save “spots” CSV file together with a folder of ROI files as input and converts them into the format required by Loon
iii. Run custom script: (*ingest_trackmate.py* available at https://github.com/visdesignlab/aardvark-util). The resulting files include metadata information (CSV, Parquet) and cell segmentation (GeoJSON).
b. Deploy Loon

i. Follow instructions on Loon Documentation: https://vdl.sci.utah.edu/loonar/docs/getting-started-with-loon/quickstart
ii. Download Docker https://www.docker.com and make an account
iii. Find and pull the latest Loon image from visdesignlab/loon
iv. Run the image and link to the directory of your data. Use volumes and environment variables specified in Loon documentation.
v. Once the Docker containers have started, visit http://localhost/ in any browser and you will be directed to Loon to interact with your data.
vi. Troubleshooting tips and videos about Loon are available on the Loon website

### Flow cytometry

Cells were harvested using 0.05% trypsin (#25300054, Thermo Fisher Scientific) and neutralized with an equal volume of cell culture media. The cell suspension was centrifuged at 500 xg for 5 min and resuspended in cold FACS buffer (0.1% BSA, 25mM HEPES, in HBSS) with or without DAPI (30ng/ml). Cell suspension was kept on ice and passed through a 35 μm cell strainer. Flow cytometry was performed using a BD Fortessa analyzer (BD, United States). SSC analysis was conducted on single cells by gating using a four-step gating strategy to exclude debris (SSC-A vs FSC-A), doublets (FSC-A vs FSC-H followed by SSC-A vs SSC-H) and dead cells (Dapi negative by gating via FSC-H vs Pacific Blue-A).

### Immunofluorescence for HMB45 and Fontana Mason Staining

MNT-1 cells were plated on a 24-well tissue culture dish. PTU was withdrawn and replaced with PTU-free media. After 5 days of culture in regular growth medium, cells were fixed with 4% paraformaldehyde (Electron Microscopy Sciences, 15710) diluted in PBS for 15 minutes at room temperature, washed 3 times in PBS, and stored at 4 °C until staining.

To detect pMEL17 positive early melanosomes, we used immunofluorescence of HMB45 (Abcam, ab787, Anti-Melanoma gp100). Cells were permeabilized for 10 minutes in 0.25% Triton in PBS, washed 3 times in PBS, blocked 30 minutes (1% BSA, 22.52 mg/ml glycine, in PBS with 0.1% Tween 20 (PBS-T)) and incubated overnight at 1:200 in antibody dilution buffer (1% BSA in PBS-T) at 4 °C. The following day, the cells were washed 3x in PBS and incubated with secondary antibody conjugated with Alexa Fluor 594 (1:1000, Invitrogen, A21203) in antibody dilution buffer and counterstained with Hoechst 33342 (10ug/ml, ThermoFisher, 62249). Cells were imaged on a Nikon Eclipse Ti2 Widefield microscope equipped with a X-cite Xylis (XT720S) light source and imaged using a 0.45 NA CF160 Super Plan Fluor PH1 ADM ELWD 20x objective. Images for mCherry (Ex: 560/40 nm, Em: 635/60 nm) and nuclear Hoescht (Ex: 375/28nm, Em: 435-485 nm) were captured with a sCMOS pco.panda camera and processed in ImageJ-FIJI (Schindelin et al., 2012).

To stain melanin, cells were processed using Fontana-Masson Stain Kit (Abcam, ab150669), following the manufactures protocol and imaged on the Nikon Eclipse Ti2 Widefield microscope using a NA 0.45 CF160 Plan Fluor PH1 DLL 10x objective with the phase condenser open in the brightfield configuration.

### Data and Code Availability

Link to interactive data hosted by Loon will be added to manuscript upon acceptance. Code will be deposited via GitHub and link with DOI will be added to manuscript upon acceptance.

## Results

### Measurement of pigmentation-associated cell phenotypes

To visualize pigmentation-associated light interactions, we developed the QUILPEN imaging pipeline using a custom-built LED-based microscope that has the flexibility to capture live cell images of light that is absorbed, phase-shifted, or scattered depending on the illumination pattern of a programmable red (624 nm) LED array used for trans-illumination (**Fig. 1a,b** and **Supplemental Fig. 1C)** (Moustafa et al., 2023). This microscope enables high-throughput imaging across multiple regions with high temporal resolution. Its integration within a standard tissue culture incubator and use of a low-phototoxicity light source allow for prolonged imaging of living samples without compromising cell viability (Polanco et al., 2022). First, four images are captured in sequence under illumination with half-circle patterns on the LED-array for reconstruction of the phase shift as light travels through a specimen by differential phase contrast (DPC) (Tian & Waller, 2015) (**Fig. 1A, C-D**). This provides a spatial map of cell dry mass (Zangle & Teitell, 2014). The resulting QPI data permit measurement of morphological features such as shape or area, identify divisions, and provide a single-cell measurement of growth rate (Nguyen et al., 2022; Popescu et al., 2008). Second, absorption, or the light that is not captured in DPC imaging because it is absorbed by the specimen, is recovered during DPC post-processing and used to generate a separate image (**Fig. 1E-F**) ((Tian & Waller, 2015), see methods). Finally, scattering information is captured using quadrant darkfield (QDF), a form of darkfield microscopy that exploits directional illumination from outside the numerical aperture, followed by mathematical removal of features that exhibit directionally biased scattering ((Moustafa et al., 2024); **Fig. 1G-I**). QDF enhances the detection of small dense objects within the cell while eliminating the peripheral scatter from cell edges, which is a limitation of traditional darkfield microscopy.

**Figure 1:**
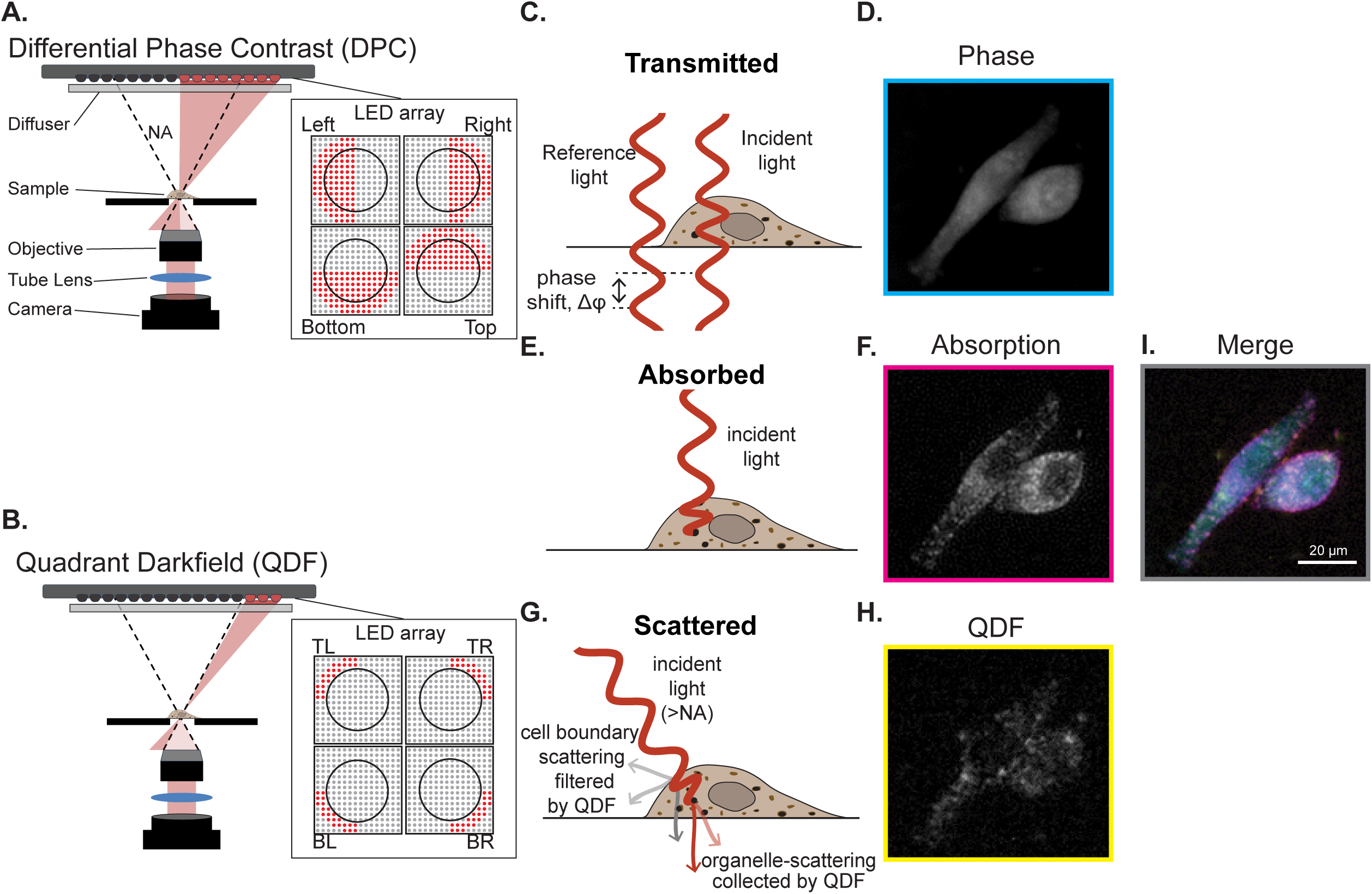
QUILPEN distinguishes cellular features that transmit, absorb, and scatter light. **A)** Schematic of an LED-array based differential phase contrast microscope (DPC) and bottom-up representation of the four LED array patterns; red indicates the LED is “on” and gray indicates “off”; black circles represent numerical aperture of the microscope objective lens. **B)** Schematic of the quadrant darkfield operation of the microscope. In QDF, four LED quarter circles are illuminated outside of the numerical aperture. **C)** Diagram of how incident light, represented by the red wave, is transmitted through a sample. **D)** Example quantitative phase image (cyan) phase shift indicated by gray value: black=0 to white= *π*/2), **E)** Diagram of the light path of absorbed light through a sample. **F)** Example absorption image (Magenta). **G)** Diagram of light path for QDF, light originates from outside the numerical aperture (NA, dotted line in A,B) scatters off small dense objects into the collection cone of the camera (yellow). **H)** Example QDF image. **I)** images of MNT-1 cell merged composite image. Lookup Table (LUT) values: Phase = 0 – *π*/2, Absorption = 0 – 10,000, and QDF = 0 – 600.

Our previous study demonstrated a strong positive correlation between scatter measured by QDF, scatter measured by flow cytometry, and visible apparent pigmentation in the cell pellet of two patient-derived acral melanoma cell lines (Moustafa et al., 2024). However, melanin exhibits both light-scattering and light-absorbing properties, and the ultrastructure of melanosomes further contributes to scattering (Cho et al., 2017; Peles & Simon, 2010). Our previous study did not resolve whether the QDF signal arises from scattering by melanin, melanosome architecture, or both. To address this, we used QUILPEN to measure additionally measure light absorption, indicative of melanin, and compared it to scatter as measured by QDF to evaluate whether differences in absorption track with QDF signal in our two previously published acral melanoma cell lines, the darkly pigmented AM084 and the lightly pigmented AM021 ((Moustafa et al., 2024; Smith et al., 2024); (**Fig 2A**)).

**Figure 2:**
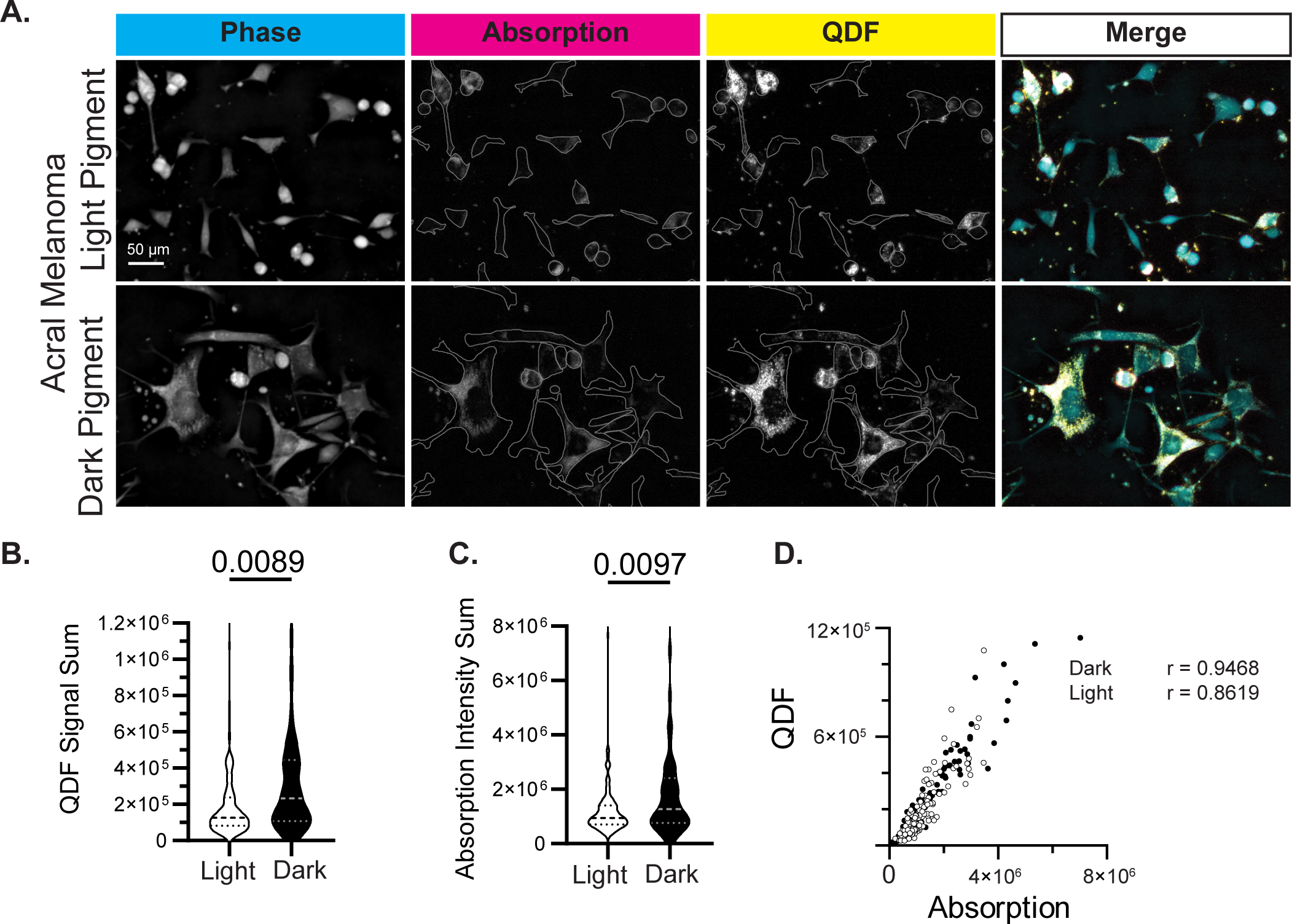
Light scattering and absorption is correlated in primary acral melanoma cells. **A**) Light (ASM21) and dark (AM84) pigmented low-passage acral melanoma cell lines imaged by QUILPEN. **B-D)** Measurements of signal sum in QDF and absorption channel. Each point represents one cell, plot represents data combined from 3 fields of view for each condition, N = 149 lightly pigmented and 98 darkly pigmented cells; Statistics: (B,C) Unpaired t test with Welch’s correction test. (D) QDF signal sum is similarly correlated with absorption signal sum for both cell types; two-tailed Spearman’s r correlation, p < 0.0001 for light and dark pigmented cells. LUT values: Phase = 0 – *π*/2, Absorption = 0 – 10,000, and QDF = 0 – 1000.

Consistent with our previous study, the highly pigmented cell line had higher scatter, as measured by QDF, and as expected, exhibited higher light absorption (**Fig. 2B,C**). Scatter (QDF) and absorption were positively correlated in both the lightly and highly pigmented lines (**Fig 2D**). This linear relationship suggests that entities present in the cells may both scatter and absorb light, or that objects that scatter light are present in the cell along with objects that absorb light. However, the mechanisms underlying the pigmentation phenotypes of these limited passage patient-derived acral cell lines remain poorly characterized. As a result, we were unable to resolve the specific contributions of pigment content versus organelle structure to QDF signal using these data alone.

### QUILPEN decouples scattered and absorbed light to distinguish cellular features by their optical properties

We turned to a well-characterized experimental model where we could measure pigment-associated changes with QUILPEN. MNT-1, a highly pigmented cell line, and SK-MEL-28, a non-pigmented cell line, are both well described in terms of pigment and melanosome content (Chen et al., 2009; Hoashi et al., 2005). As expected, MNT-1s appeared darkly pigmented when pelleted and exhibited significantly higher absorption relative to SK-MEL-28s (**Fig. 3A-C**). Similarly, we observed an overall stronger QDF signal in the MNT-1s compared to the SK-MEL-28s (**Fig. 3D**). However, the difference was smaller than what we observed in the absorption measurements, and the SK-MEL-28 population exhibited a broader QDF signal distribution than MNT-1s (**Fig. 3C-D**). While both cell types differ markedly in pigment content, they each contain melanosomes. MNT-1 cells predominantly harbor mature, melanin-rich stage III–IV melanosomes, whereas SK-MEL-28 cells contain mostly immature, stage I–II melanosomes. (Chen et al., 2009). Our QDF data suggest that SK-MEL-28 cells, despite lacking visible pigment, can exhibit total melanosome content equal to, or greater than, the total melanosome content of MNT-1 cells. Notably, the positive correlation between QDF and absorption observed in the acral melanoma lines was maintained in the MNT-1/SK-MEL-28 system. However, the proportional relationship between QDF and absorption, indicated by the slope, was different between the acral lines, the MNT-1s, and the SK-MEL-28s, (**Fig. 3E**), further suggesting that QDF reflects organelle content and is not driven by melanin-based scattering or dependent on melanosome maturity.

**Figure 3:**
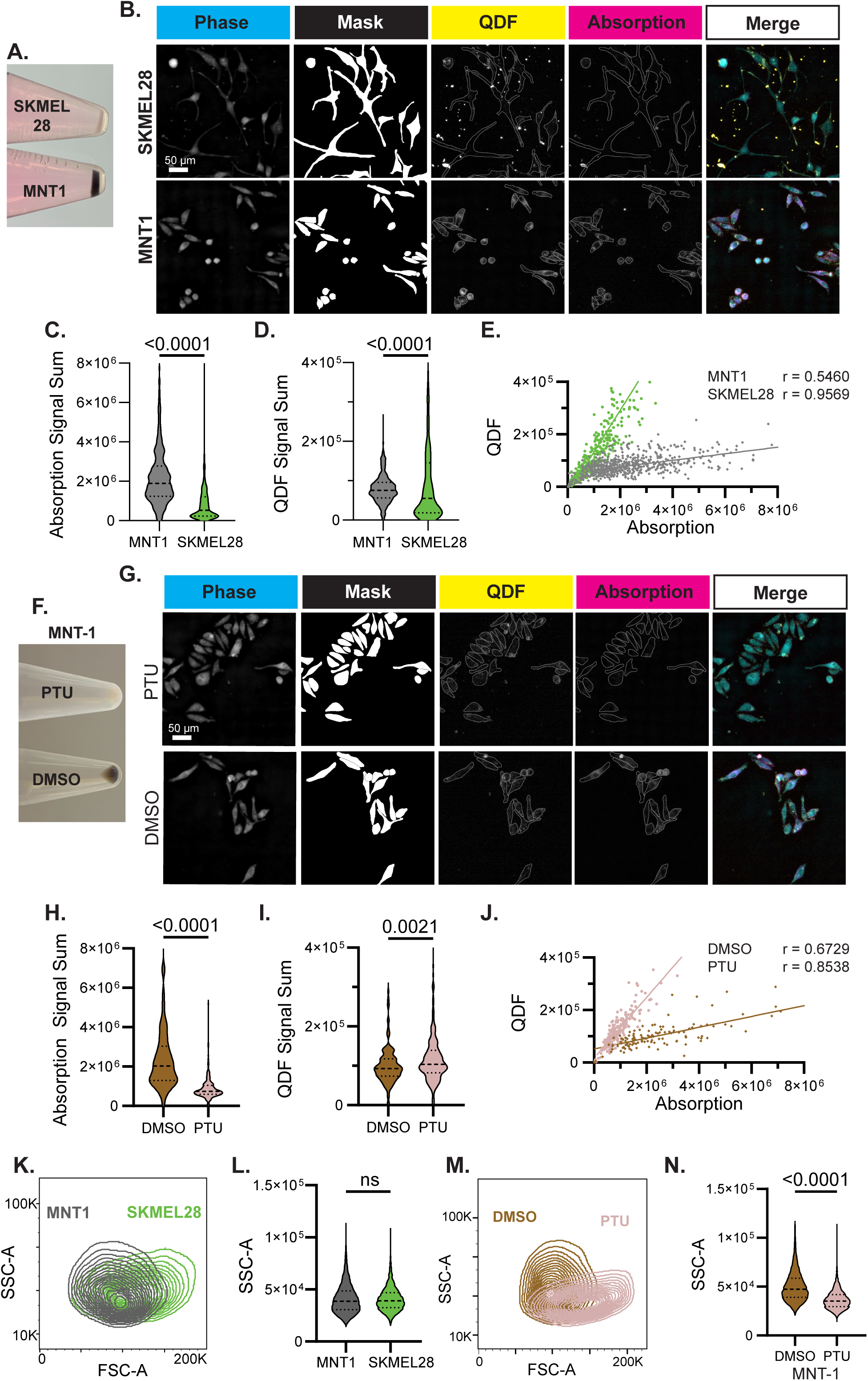
QUILPEN decouples scattered and absorbed light to distinguish cellular features by their optical properties. **A)** Cell pellets from SK-MEL-28s (non-pigmented) and MNT-1s (darkly pigmented) melanoma cell lines. **B)** Melanoma cell lines imaged by QUILPEN. **C,D)** Measurements of signal sum in Absorption and QDF channel. Each point represents one cell, plot represents data combined from 3 fields of view for each condition, N = 551 SK-MEL-28 and 941 MNT-1 cells; **E)** QDF and absorption signal sum are correlated; p < 0.0001 for both SK-MEL-28 and MNT-1 cells, but distinct between cell lines, p<0.0001. **F)** cell pellets of MNT-1 cells treated for 21 days with PTU (non-pigmented) or DMSO control (pigmented). **G)** MNT-1 cells imaged by QUILPEN after DMSO or PTU treatment. **H,I)** Measurements of signal sum in Absorption and QDF channel. Each point represents one cell, plot represents data combined from 3 fields of view for each condition, N = 132 DMSO and 264 PTU treated cells; **J)** QDF and absorption signal sum are correlated; p <0.0001 for both PTU and DMSO treatment conditions, but distinct between conditions, p < 0.0001. **K,L)** Flow cytometry measurement of granularity by side-scatter in MNT-1 and SK-MEL-28 cells. **M,N)** Side-scatter analysis of MNT-1 cells after DMSO or PTU treatment. Statistics: (C,D,H,I,L,N) Unpaired t test with Welch’s correction test, (E, J) r from two-tailed Spearman’s correlation, line from linear regression with slope comparison. LUT values: Phase = 0 – *π*/2, Absorption = 0 – 10,000, and QDF = 0 – 600.

To directly test whether QUILPEN can distinguish melanin content from organelle abundance, we used 1-phenyl-2-thiourea (PTU), a well-characterized reversible inhibitor of tyrosinase activity, to reduce pigment levels in MNT-1 cells (Chen et al., 2009). Long-term PTU treatment prevents melanin synthesis, leading to a gradual loss of pigmentation without disrupting melanosome biogenesis. This pigmentation phenotype is well documented in the MNT-1 melanoma line, where PTU-treated cells shed pigmented melanosomes over time but retain a steady-state number of melanosomes (Chen et al., 2009). We cultured MNT-1 cells in PTU or DMSO (vehicle control) for 21 days, resulting in a visibly depigmented cell pellet in the PTU-treated condition (**Fig. 3F**). PTU treated cells exhibited lower light absorption relative to DMSO control but retained similar levels of QDF signal, consistent with loss of melanin but retention of melanosomes (**Fig. 3G, H)**. Similar to the relationship between the non-pigmented SK-MEL-28s and MNT-1s, the PTU treated MNT-1s had a steeper slope, with a greater QDF signal relative to the absorption signal when compared to the DMSO treated MNT-1s (**Fig. 3J**). HMB45 staining, a marker of processed internal domain of Pmel17 (Hoashi et al., 2005), was comparable between conditions, confirming that PTU treatment did not impact melanosome content (**Supplemental Fig 2**), and further supporting the observed stability of the QDF signal.

To benchmark QUILPEN’s ability to decouple melanin content from organelle abundance, we compared our imaging-based measurements to side scatter (SSC) from flow cytometry. SSC is commonly used as a proxy for pigment content (Bajpai et al., 2023; Belote et al., 2021; Choi et al., 2012; Moustafa et al., 2024). Although SSC has been shown to correlate with melanin levels, both in our prior work and that of others, the extent to which it specifically reflects pigment, rather than melanosome number or structure, remains unclear. If SSC were primarily driven by melanin, we would expect a marked difference between MNT-1 and SK-MEL-28 cells as observed in the two acral melanoma lines (Moustafa et al., 2024). However, MNT-1 and SK-MEL-28 exhibited similar SSC profiles (**Fig. 3K, L**). In contrast, if SSC were primarily driven by organelle scatter, we would expect no difference between MNT-1 cells treated with PTU and DMSO. However, PTU and DMSO treated MNT-1 cells had significantly different SSC profiles (**Fig. 3M,N**). Together, these data indicate that SSC signal does not discriminate between granularity and melanin content, whereas QUILPEN uniquely deconvolutes signals from melanin content versus organelle abundance.

### Repigmentation is captured by absorption imaging, but not QDF

To determine whether QUILPEN can measure changes in melanin content, we performed live imaging of MNT-1s treated with PTU or DMSO control after the removal of treatment (**Fig. 4A,B**). Cell division occurred over the full 5-day experiment, with the expected exponential increase in cell number (**Supplementary Fig. 3A**). The pigment levels of PTU treated cells were restored and the absorption signal converged with the pigment levels of DMSO treated cells within the imaging time course (**Fig 4B, C**), consistent with the return of pigment that could be observed by Fontana Mason staining (**Fig. 4D**).

**Figure 4:**
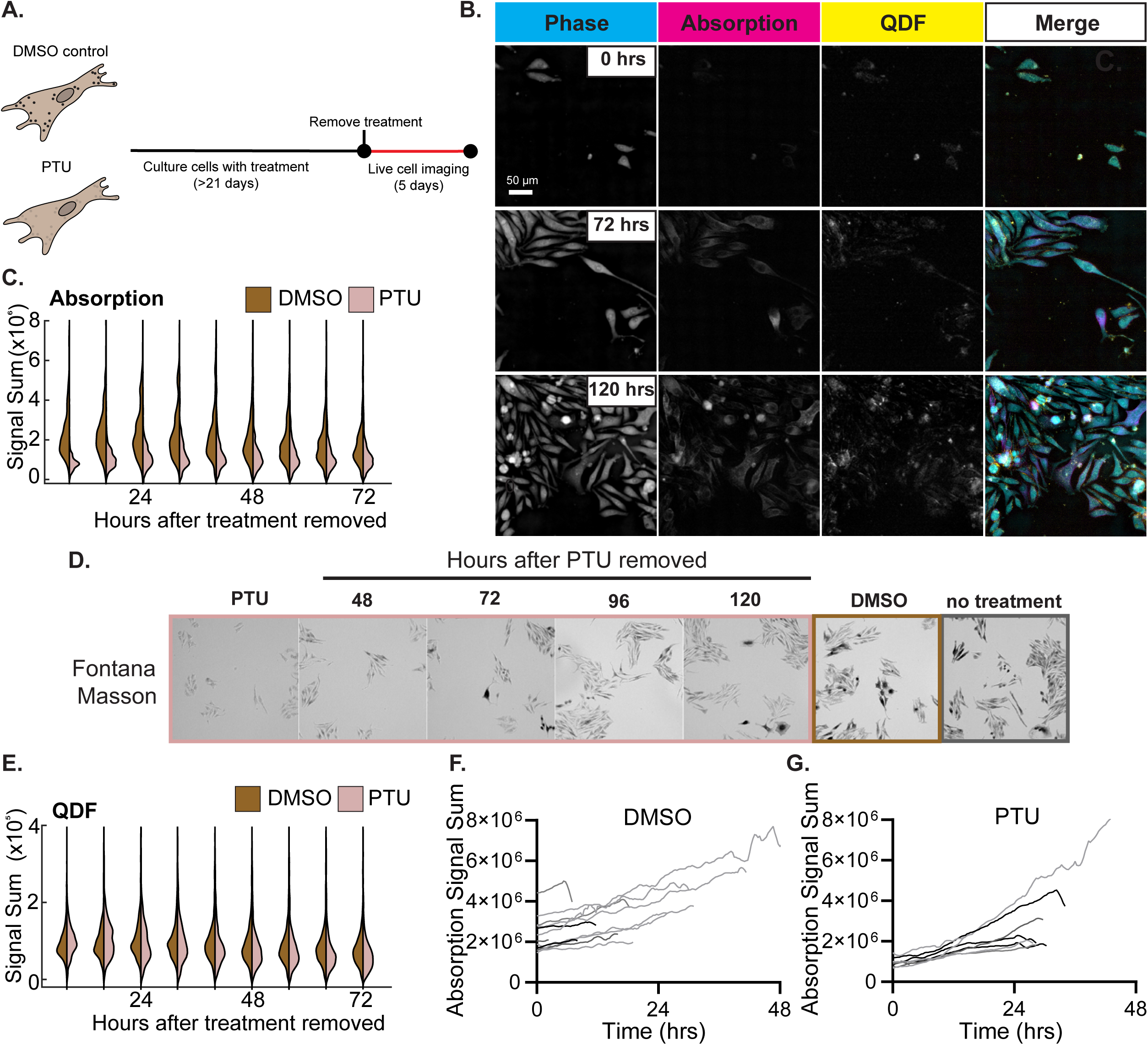
Absorption signal captures the heterogenous dynamics of repigmentation after PTU removal. **A)** Schematic of PTU treatment and imaging workflow. **B)** MNT-1 cells imaged by QUILPEN at the start of experiment, 72, and 120 hours after PTU removal. **C)** Absorption signal sum grouped in 8-hour time bins for 72 hours. For cell number in each bin see Supplementary Fig. 3A. **D)** Fontana Masson staining for melanin repigmentation on cells fixed at indicated timepoint after PTU removal. **E)** QDF signal sum, same cells measured as in C. **F,G)** Signal sum from tracking of first-generation cells after DMSO and PTU removal. N = 70 for DMSO and N= 87 for PTU treated cells. Statistics: (C,E) Unpaired t test with Welch’s correction test. LUT values: Phase = 0 - *π*/2, Absorption = 0 – 10,000, and QDF = 0 – 600.

Notably, the QDF signal was stable and similar between the DMSO and PTU treated cells after the removal of treatment indicating a steady state of melanosome content irrespective of melanin content (**Fig.4E, Supplementary Fig. 3 B,C**).

Under steady-state conditions, untreated and DMSO-treated MNT-1 cells exhibited a wide range of absorption signal, revealing intrinsic heterogeneity in melanin content (**Fig. 3C, H**; **Fig. 4C, D**).

Remarkably, this distribution of absorption levels was re-established during repigmentation following PTU removal (**Fig. 4C, D**), suggesting that cells restore not only total pigment but also their relative pigmentation states. To investigate this further, we tracked individual cells over time, focusing on the generation of cells present at the time of treatment removal. DMSO-treated cells showed a steady increase in absorption over the life cycle of the cell (**Fig. 4F**). In contrast, the PTU-treated population displayed substantial variability in the kinetics of pigment recovery (**Fig. 4G**), revealing cell-intrinsic differences in the rate and extent of melanin restoration. The QDF signal had fluctuations throughout the life cycle of the cell and was variable within the population for both PTU and DMSO-treated cells. However, no directional change was evident, suggesting organelle content was maintained during cell growth (Supplementary Fig. 3B, C). In summary, the temporal measurements of absorption and QDF in bulk and individual cells showed heterogeneity in both the amount and rate of repigmentation, while organelle content remained stable.

### Cell lineages have varied inheritance of pigment related features

Given the variability of pigmentation recovery, we next asked whether the pigment heterogeneity arose from clonal expansion of highly pigmented cells or from differences in pigmentation dynamics within lineages. We first used the QPI component of QUILPEN to segment and track single cells. Then, using Loon, an interactive data visualization tool (Lange et al., 2022, 2025), we built lineage trees and overlaid absorption and scattering data to examine inheritance patterns (**Supplemental Fig. 4,** see data availability statement). Tracking changes in a single lineage over multiple cell divisions highlights the heterogeneity in cell cycle length and pigment inheritance. Cells underwent one to five division cycles over the 5-day experiment (Fig. 5A, Supplemental Fig. 4). In the DMSO-withdrawn condition, a single lineage had increasing pigment throughout cell growth and variation in pigment inheritance after division. Interestingly, there was a trend of decrease in pigment content in daughter cells following division after 72 hours. (**Supplemental Fig. 4B, Fig 4C**). Cells in the PTU-withdrawal condition showed varied rates of absorption increase during the cell growth phase and inheritance patterns. In one representative example (Lineage 1), a parent cell divided only once but exhibited rapid absorption recovery relative to neighboring cells (**Fig 5A, B**). The division produced two daughter cells with similar absorption levels, indicating symmetric inheritance of pigment content. In contrast, a neighboring cell (Lineage 2) underwent five divisions and displayed both symmetric and asymmetric inheritance of absorption between daughter cells (**Fig. 5 A, C**). In both lineages, daughter cells continued to show variable rates of absorption increase after division, indicating that pigment recovery is not solely determined at the time of inheritance. QDF signal also increased over time in both lineages, with a greater relative increase observed in Lineage 1. (**Fig. 5 D, E**). However, the magnitude of QDF change was smaller than that of absorption. In Lineage 2, QDF also increased gradually but to a lesser extent. In both lineages, daughter cells exhibited similar QDF levels after division and comparable rates of increase, indicating symmetric inheritance of scatter-associated features (**Fig. 5D,E**). These findings suggest that organelle content is distributed equally between daughter cells, while pigment-loaded melanosomes are unequally inherited, resulting in enrichment of pigment in one sibling. The variable rates of absorption increase further suggest that the capacity for pigment synthesis may also be asymmetrically inherited, independent of melanosome number.

**Figure 5:**
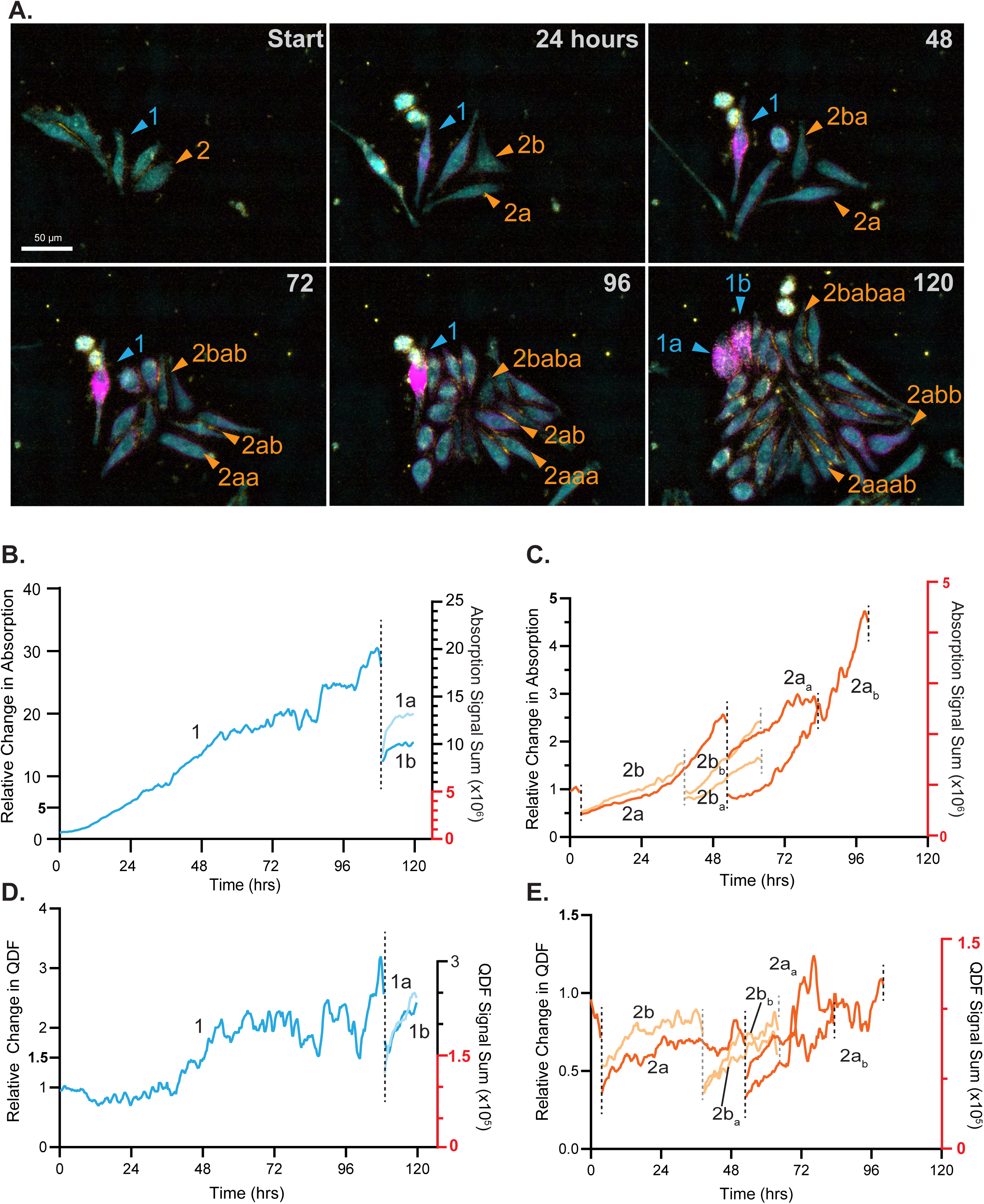
Cell lineages have varied inheritance of pigmentation-related features. **A)** QUILPEN merged images of MNT-1 cells acquired for 120 hours after PTU removal. Colored arrowheads and labels denote lineage with “a” and “b” appended for each generation to indicate siblings. **B,C)** Absorption over time for lineage 1 (blue) and lineage 2 (orange). Plots include up to two generations after the parental cell at the start of experiment. Left y-axis indicates the relative change in absorption from initial value, right y-axis indicates absorption signal sum. Dotted vertical lines indicate cell division and, where applicable, connect a dividing cell to its offspring. Red segment of right y-axes is included to draw attention to difference in axes range between the two lineages. **D,E)** QDF over time following the same color and labeling conventions as B and C.

## Discussion

We developed QUILPEN, a live-cell imaging and analysis platform that decouples quantification of melanin content from melanosome abundance in individual cells. Our approach enables dynamic, label-free tracking of pigment-associated features with single-cell resolution across time. By independently quantifying absorbed and scattered light, QUILPEN distinguishes melanin content from intracellular granularity and morphological complexity (**Fig. 1**). Applying this approach to established melanoma cell lines, we identified substantial heterogeneity in both melanosome content and pigmentation, even among genetically identical cells (**Fig. 3**).

Using SK-MEL-28 and MNT-1 cells, two melanoma cell lines that differ markedly in melanin content but not necessarily in melanosome number, we found that the relationship between the scattering and absorption signals were distinct (**Fig. 3E**). This separation was not apparent in conventional flow cytometry, where side scatter integrates contributions from both melanin and organelle content (**Fig. 3K,L**). Recovery of pharmacologic inhibition of melanin synthesis by PTU further demonstrated that pigment levels recovered while melanosome abundance remained stable (**Fig. 4; Supplemental Fig. 3, C**), reinforcing the distinction between pigment synthesis and organelle biogenesis. Thus, QUILPEN exploits the optical properties of melanosomes to uniquely enable a level of optical specificity and biological interpretability that is not achievable with current label-free approaches.

Longitudinal imaging of pigment recovery following PTU removal revealed striking variability in the kinetics of melanin restoration within a clonal MNT-1 population (**Fig. 5**). Lineage tracing showed that these differences were not only cell-intrinsic but also heritable, with asymmetric transmission of pigmentation recovery phenotypes to daughter cells. In the context of melanoma, this type of heterogeneity could have important consequences. Pigmentation affects oxidative stress buffering, immune visibility, and drug sensitivity - features that influence tumor progression and therapeutic resistance. While clinical observations and bulk transcriptomic analyses have long noted pigmentation heterogeneity within tumors, direct measurements of melanin content at single-cell resolution have been lacking. Prior studies have relied on either bulk assessment of population pigmentation or single-cell transcriptomics, neither of which directly report melanin levels in individual cells. The longitudinal imaging of the DMSO-withdrawn cells also revealed a decline in the pigmentation over time in both the bulk analysis and within individual lineages. This observation suggests that changes in the culture conditions, such as nutrient concentration, pH, or cellular confluence, may influence pigmentation. This is unsurprising, as pigment production is dependent on pH of the media and pH decreases over the 5-days in the incubator, indicated by yellowing of the phenol red containing media (Ancans et al., 2001). QUILPEN now makes it possible to connect molecular, morphological, and behavioral features of melanoma cells with actual pigment content. Future work can also address the use of QPI-derived dry mass and cell morphology to assess changes in single-cell growth rates and morphology during melanogenesis.

There are several technical considerations to address in future work. One is the effect of illumination wavelength on the interpretation of absorption and scattering. In this study, we used ∼624 nm red light, which aligns well with the optical properties and size of mature melanosomes. However, different wavelengths may offer selectivity for eumelanin versus pheomelanin, may distinguish other dense intracellular structures. Expanding the spectral capabilities of QUILPEN could broaden its utility to include a wider range of pigment types and organelle classes. Notably, QUILPEN’s optical approach differs from that of flow cytometry, where side scatter is typically measured using a 405 nm laser (e.g., on a BD Fortessa). This spectral difference could contribute to the divergence in findings between QUILPEN and SSC. Another promising direction is integration with fluorescence-based reporters, which could link organelle and melanin content to gene expression, intracellular pH, metabolic state, and other molecular markers in the same cells over time. Such combined measurements would expand the scope of QUILPEN to address how melanin production intersects with transcriptional programs, differentiation state, stress responses, and disease progression. A pitfall of this work is the tedious manual correction required for accurate segmentation and tracking, which hinders high-throughput analysis. However, emerging machine learning tools are rapidly improving segmentation of quantitative phase images and enhancing multi-lineage tracking predictions, reducing the need for manual intervention.

In summary, QUILPEN provides a new platform for studying melanogenesis and melanosome biology with high temporal and spatial resolution. It enables precise dissection of pigment synthesis dynamics, inheritance, and heterogeneity in live cells, without labels or perturbations. By decoupling pigmentation from other cellular features, QUILPEN opens new opportunities to explore how melanin production interfaces with gene expression, differentiation state, stress response, and disease progression. These insights are broadly relevant not only to melanoma, but also to pigmentary disorders, developmental biology, and the functional diversification of melanocytic cells across tissues.

## Resources

We thank the Flow Cytometry Shared Resource at The Ohio State University Comprehensive Cancer Center, Columbus, OH for providing access to flow cytometry instrumentation.

## Author Contributions

Conceptualization: R.G.Z, R.L.J.-T., T.A.Z., R.L.B; Methodology: R.G.Z, S.A, T.E.M, T.A.Z, R.L.B; Scripting: R.G.Z, S.A, T.E.M, T.A.Z; Microscope updates and maintenance: R.G.Z, S.A, T.E.M.; Formal Analysis: R.G.Z, R.L.B; Data Visualization: R.G.Z,, D.L., L.S., R.L.B; Investigation: R.G.Z, S.A, T.A.Z, R.L.B, Writing – original draft: R.G.Z and R.L.B; Writing – Review and editing: R.G.Z., R.L.J.-T, T.A.Z, R.L.B.; Supervision: R.L.J.-T., T.A.Z, R.L.B; Funding acquisition: A.L, R.L.J.-T., T.A.Z, R.L.B,

## Data Availability

Data supporting the findings of this study are available at {Website links with appropriate DOI will be shared upon acceptance}.

## Code Availability

Custom code is available at GitHub {GitHub link with DOI will be shared upon acceptance}

