## Supplemental Figures for "QUILPEN provides independent and label-free single-cell quantification of pigmentation dynamics and organelle content"

### Supplementary Figure 1

**A. Protocol A: Three-Channel Image Acquisition and Processing**

1. Microscope construction

2. Acquire images

3. Matlab output to TIFF images

↓

**B. Protocol B: Image Analysis for Feature and Lineage Tracking**

**1. High fidelity analysis (Fig. 2, 3, 4F-G, 5)**

1a. Export selected frame(s) using ImageJ

↓

1b. Segment Cells with Cellpose

↙ ↘

1c. Record measurements (Single frame, Fig. 2-3)

1d. Trackmate with label image (Time course, Fig. 4F-G,5)

**2. Bulk cell analysis (Fig. 4 C,E)**

2a. Prepare 3-channel image stack

↓

2b. Segment using TrackMate-Cellpose

↓

**3. Track post-processing (Fig. 4)**

↓

3a. convert data for plotting

↓

3b. Prepare plots
(Fig. 4 C,E - Split Violin)
(Fig. 4, F-G - Line Plot)

**4. TrackMate to Loon (Fig. 5)**

4a. Reformat TrackMate output for Loon

↓

4b. Deploy Loon

**C.**

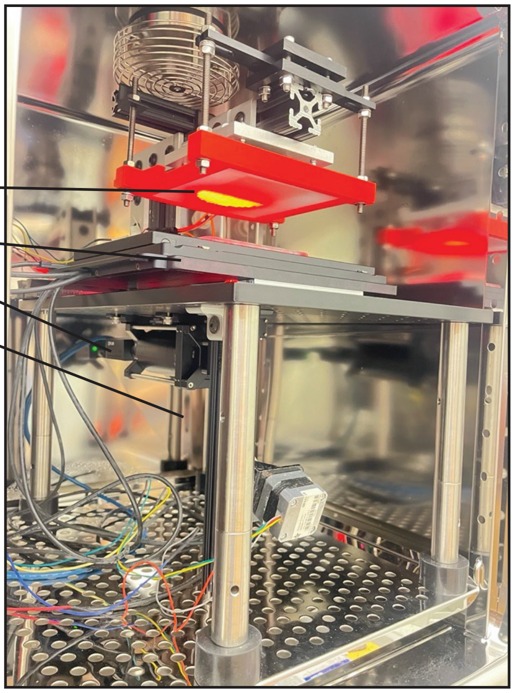

### Supplementary Figure 2

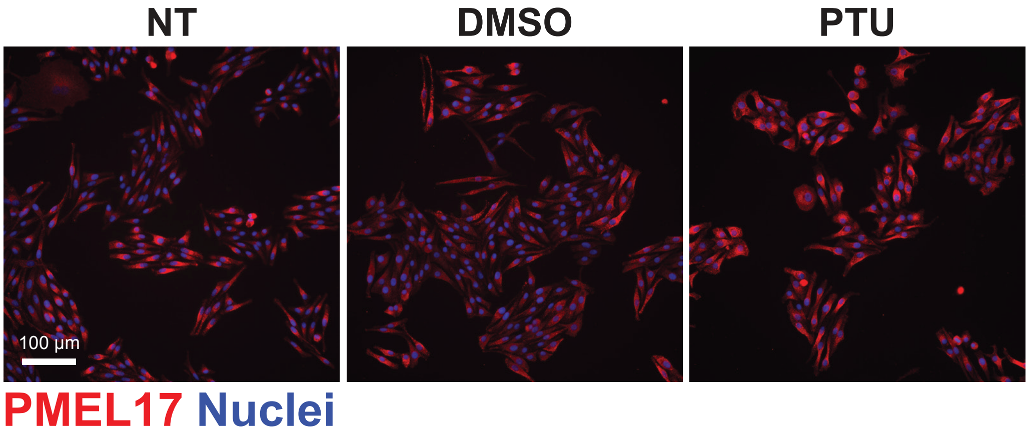

**Supplementary Figure 3**

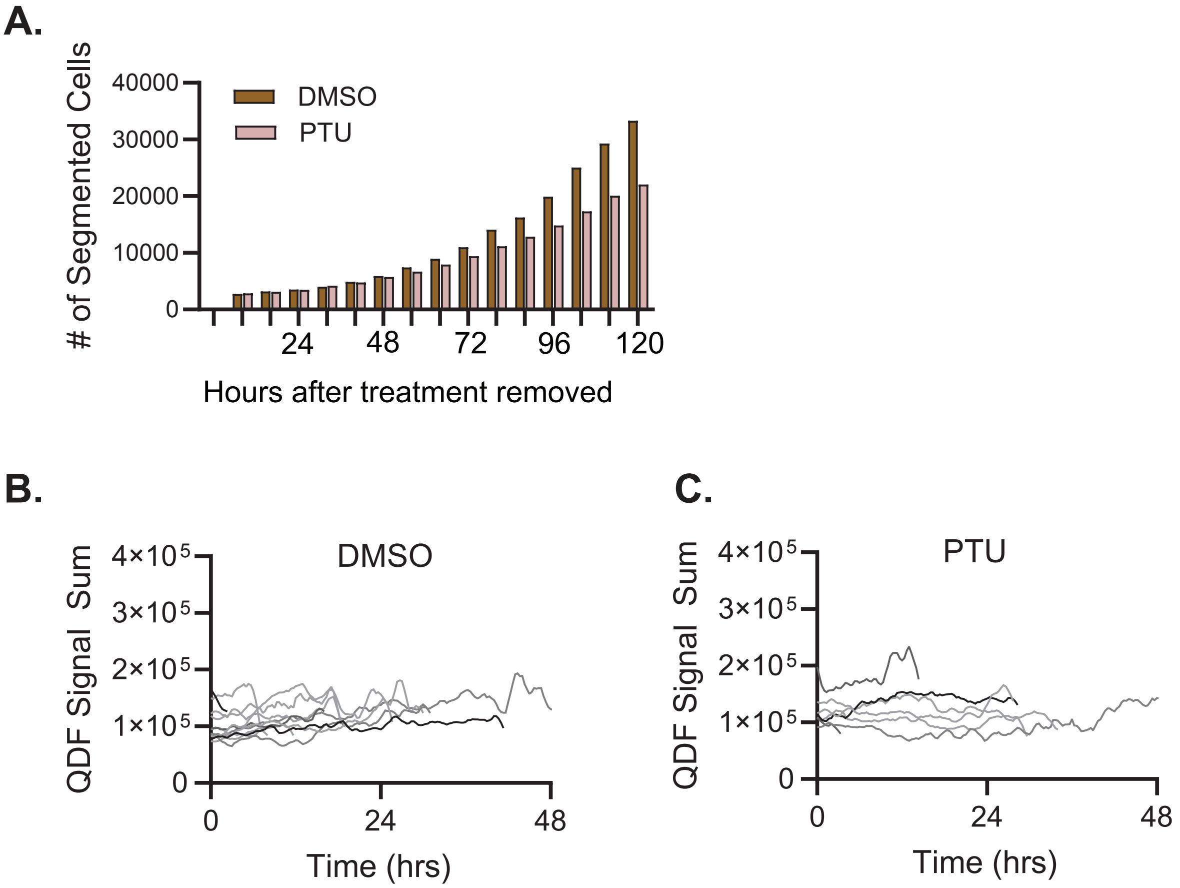

### Supplementary Figure 4

**A.** Looneage

DMSO, Single Lineage, Displaying 4 of 6 generations

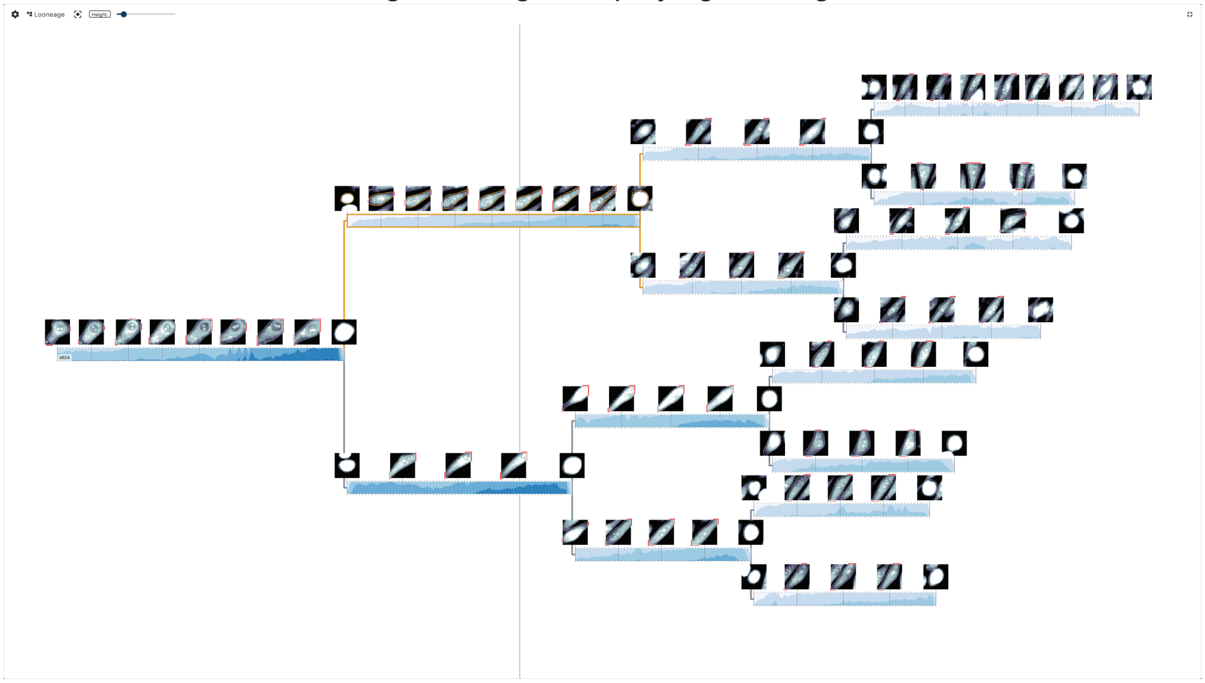

**B.** Line Chart

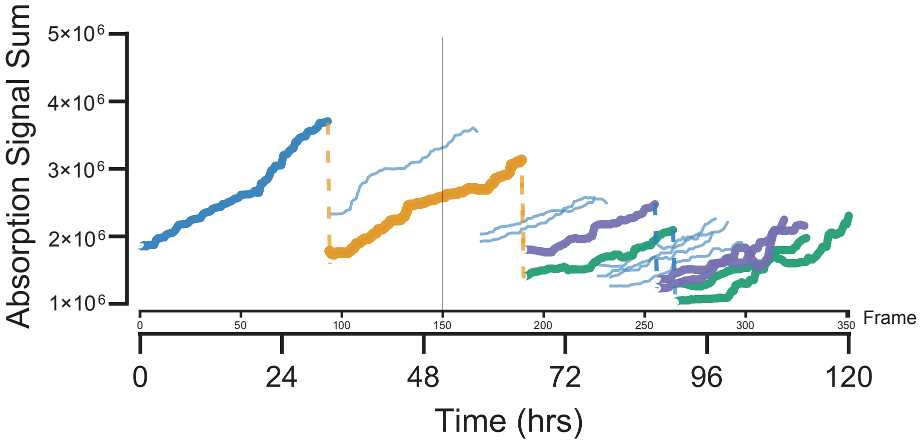


**C.** Images

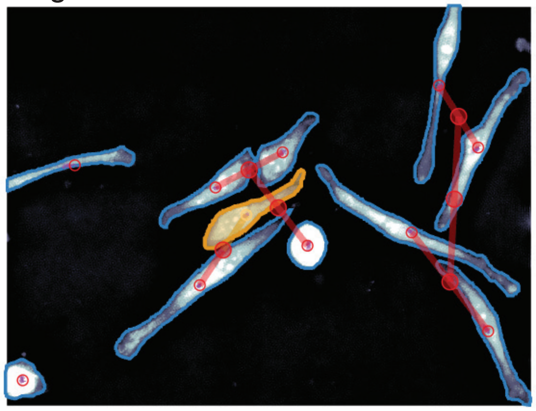
