## Supplemental Figure Legends for "QUILPEN provides independent and label-free single-cell quantification of pigmentation dynamics and organelle content"

*Supplemental Figure 1: Optical System*

**A)** Workflow of three-channel image acquisition and processing to accompany Protocol A. **B)** Workflow of Image Analysis for feature and lineage tracking to accompany Protocol B. **C)** Photograph of the custom-built LED-based microscope in a cell culture incubator to maintain CO_2_ and temperature control for the 5-day experiment.

*Supplemental Figure 2: MNT-1 cells have similar melanosome staining across all PTU treatment conditions.*

Early melanosomes (red) in non-treated (NT), DMSO-withdrawn, and PTU-withdrawn MNT-1 melanoma cells by immunofluorescence using HMB45 (mCherry) and nuclei (Hoechst).

*Supplemental Figure 3: QDF signal is heterogenous but steady relative to absorption in DMSO and PTU treatment conditions*

**A)** Histogram of cell count per bin for Fig 4 C, by condition. **B,C)** QDF signal sum after Cellpose segmentation and TrackMate based tracking of cells after DMSO and PTU withdrawal in manually verified parent tracks.

*Supplemental Figure 4: Loon interactive visualization of cell tracking*

**A)** Loon tree-style interface displaying four of six total generations of a cell lineage of MNT-1 cells after DMSO treatment. The start time of the experiment is when DMSO was removed. Thumbnail images represent selected frames throughout the dataset. Initial and final images in a track show rounded up cells during mitosis and elongated cells during the growth phase in the middle of the track. The blue colorbar represents cell area, with a darker color indicating higher values. **B)** Line-chart style visualization of the same four generations as in (A) with the y-axis indicating the attribute absorption signal sum. **C)** Image of selected cell (yellow) at the timepoint indicated by the vertical line (150 frames, 50 hours). Cell outlines are in blue and red lines indicate lineage traces linking sibling cells. Yellow highlighted tracks in A and B match the selected yellow cell in C.
